# Novel strains of *Campylobacter* cause diarrheal outbreak in Rhesus macaques (*Macaca mulatta*) of Kathmandu Valley

**DOI:** 10.1101/2022.06.20.496768

**Authors:** Rajindra Napit, Prajwol Manandhar, Ajit Poudel, Pragun G. Rajbhandari, Sarah Watson, Sapana Shakya, Saman M. Pradhan, Ajay N. Sharma, Ashok Chaudhary, Christine K. Johnson, Jonna K. Mazet, Dibesh Karmacharya

**Author notes:** These authors contributed equally to this work (Rajindra Napit, Pragun G. Rajbhandari).

## Abstract

*Campylobacter spp*. is often underreported and underrated bacteria that present real health risks to both humans and animals, including non-human primates. It is a commensal microorganism of gastrointestinal tract known to cause gastroenteritis in humans. Commonly found in many wild animals including non-human primates (monkeys-Rhesus macaques) these pathogens are known to be a common cause of diarrhea in humans in many parts of developing and under developed countries.

Rhesus macaques from the two holy sites in Kathmandu (Pashupati and Swoyambhu) were included in this cross-sectional study. Opportunistic diarrheal samples of monkeys were analyzed to detect and characterize the pathogen using 16S rRNA-based PCR screening, followed by DNA sequencing and phylogenetic analysis.

Out of a total 67 collected diarrheal samples, *Campylobacter spp*. were detected in the majority of the samples (n=64; 96%). DNA sequences of the amplified PCR products were successfully obtained from 13 samples. Phylogenetic analysis identified *Candidatus Campylobacter infans* (n=10, Kimura-2 parameter (K2P) pairwise distance values of 0.002287). Remaining three sequences might potentially belong to a novel Campylobacter species/sub-species-closely relating to known species of *C. helviticus* (K2P pairwise distance of 0.0267). Both *Candidatus Campylobacter infans* and *C. helvitucus* are known to infect humans and animals. Additionally, we also detected the bacteria in water and soil samples from the sites. *Campylobacter spp*. caused the 2018 diarrhea outbreak in Rhesus macaques in the Kathmandu valley. *Campylobacter* might be one of the important contributing pathogens in diarrheal outbreaks-both in humans and animals (monkeys) in Nepal. Due to close interactions of these animals with humans and other animals, One Health approach might be the most effective way to prevent and mitigate the threat posed by this pathogen.

## Introduction

*Campylobacter spp*. is a zoonotic pathogen that is found in the gut flora of many species ranging from domesticated to wild animals-both free roaming and in captivity. It is capable of infecting humans as well as non-human primates (NHP) causing mild to severe gastrointestinal problems [1–3]. Emergence of antibiotic-resistant strains of *Campylobacter* is a major public health concern as animals carrying the bacteria pose a significant risk to humans via contamination of water sources, food, or through repeated interactions (physical contacts) [2–4]. In Nepal, *Campylobacter* is one of the leading causes of food-borne infections; and antibiotic resistant strains of the bacteria have also been reported in poultry slaughter houses throughout Nepal [5–8]. Most of the studies conducted in the country have mostly been limited to food-borne pathogenesis of *Campylobacter* [5–9]. Although, zoonotic spillover is highly prevalent from interactions with animals either carrying or infected with *Campylobacter*, very limited studies have been conducted in Nepal [6]. Limited publications are available documenting spillover from domesticated dogs *Canis lupus familiaris* [10] and from livestock to farmers [11]. However, no such studies have been published highlighting detection and possible spillover of *Campylobacter* from NHP such as Rhesus macaques *(Macaca mullata, commonly known as monkeys)* to humans in Nepal, even though, risk of such exposure and disease transmission have been widely documented [2–4].

*Campylobacter spp*. have been found in both captive and free-roaming monkeys causing diarrhea and severe enterocolitis [2,12,13]. Due to NHP’s high genomic similarities and close evolutionary relationships to humans, and similar gut flora detected in developing countries, the risk of zoonotic transfer of pathogenic strains of bacteria like *Campylobacter* is highly probable [14,15]. Previous studies have detected presence of the bacteria in captive NHPs in the United States [16], New Zealand [17], and in Kenya [13]. Presence of pathogenic strains of the bacteria in healthy and asymptomatic monkeys[1] poses even higher risk of zoonotic spillover through direct physical contact or indirectly as a potential source of environmental contamination by fecal matter [18].

In Kathmandu, monkeys inhabit few of the major holy sites including Pashupati and Swoyambhu. These sites, are surrounded by dense urban human population and have significant human-wildlife (monkey) interactions. Following report of diarrhea outbreak (2018) in monkeys of these two sites, we carried out an opportunistic cross-sectional research- specifically designed to detect and characterize *Campylobacter* infections.

## Materials and Methods

### Ethical Statement

The research was conducted as a supplementary study to the PREDICT project which focused on understanding emerging diseases in urban-wildlife interfaces. All the permits and ethical clearance were obtained before the study. The non-invasive sampling and analysis were covered by the permit obtained from the Nepal Health Research Council (NHRC, Ref.no. 224).

### Study Design and Site Selection

A cross-sectional study was conducted during active diarrheal outbreaks at two Rhesus macaque inhabiting sites (Pashupati and Swoyambhu) in Kathmandu (Nepal) in 2018 (June -July) [19]. The sampling sites were chosen according to the confined habitat, with a focus on areas with frequent monkey-human interactions. These two holy (temple) sites of in the Kathmandu have many free-roaming monkeys and are frequently visited by people for sightseeing and religious purposes. Cattle, dogs, chickens and other birds are also present in abundance in neighboring areas of the temples.

Swoyambhunath temple, one of the oldest Buddhist holy sites in the region, is situated on top of a hillock in the northwest of the Kathmandu Valley. Also known as the “Monkey Temple”, the area surrounding Swoyambhunath (with area of 2.5 square kilometer) is home to one of the largest populations of free-roaming macaques in the region. The site hosts an estimated population of 400 monkeys [20]. Pashupatinath temple is one of the most important and popular holy Hindu sites in Nepal. This site is home to a population of 300 monkeys [20,21], which reside in nearby patches of forests (Bankali, Bhandarkhal and Mrigasthali) surrounding the temple premises [22].

Both sites were divided into 5 transects, and 5 field teams were mobilized to systematically comb through each transect and collect only diarrheal (loose) fecal samples from monkeys. Feces were collected using sterile swabs in a tube containing phosphate-buffered saline (PBS). A portion of the feces were also collected in silica gel tubes as replicate. Additionally, adjacent soil and water samples from a drinking water source (pond) were also collected to check for any possible environmental contamination. The samples were immediately transported in cold-chain to our lab in Kathmandu and stored at -20°C freezer for further processing. A total of 67 opportunistic diarrheal fecal samples and surrounding soil samples (n=11) were collected from both the sites. Some water samples (n=5, river and drinking water sources) were also collected.

### Campylobacter detection and characterization

#### Molecular screening and 16S rRNA gene sequencing

Bacterial DNA was extracted from collected samples using Bacterial DNA extraction kit (Zymo Research, USA) according to the manufacturer’s instructions, and were stored at -20°C. Screening for *Campylobacter* was carried out by targeting the ∼ 800 bp fragment of the 16S rRNA gene using *Campylobacter* genus specific PCR primer sets C412F and C1288R [23]. PCR amplification was done in 25µl volume containing reaction buffer, 0.2nmol primers, Taq polymerase and 2ul template. The cycling condition for the PCR-initial denaturation at 95°C for 4 minutes, 35 cycles of denaturation at 95°C for 30 sec, annealing at 55°C for 30 sec and extension at 72°C for 30 sec, final extension 72°C for 10 min and hold at 4°C. The PCR products were separated on 1.5% agarose gel electrophoresis.

8µl amplified PCR product was cleaned using 2µl of ExoSAP-IT™ kit (Thermofisher, Catalog No. 78200.200.UL). The reaction mixture was then incubated for 30 minutes at 55°C to get rid of excess primers, followed by 85°C for 10 minutes for reaction deactivation. The purified PCR products were then sequenced on an ABI thermocycler using BigDye^™^Terminator V3.1 Cycle Sequencing Kit (Catalog No. 4337455). Excess salts and dye terminators were removed using BigDye® XTerminator™ Purification Kit (Catalog No.4376486). The samples were then analyzed on ABI 310 Genetic Analyzer.

### Phylogenetic Analysis

The phylogenetic analysis was performed using BEAST v2.6.4 of partial 16S rRNA sequence [24]. A ∼649bp (final size after quality trim) sequences from the NCBI Genbank database was obtained of all known *Campylobacter* species [25]. Along with the sequences obtained from the NCBI database, 13 sequences (Genbank acc: MZ06810 to MZ068112) obtained from this study was also included to form a dataset of 16S rRNA partial sequences (supplementary table 2). All the sequences were aligned using MUSCLE v3.8.425 [26] and was visualized in AliView v1.27 [27]. Model test for BEAST analysis was performed using Bmodel test v1.2.1 [28] for substitution model for 10 million iterations. The phylogenetic tree was prepared on BEAST v2.6.4 with HKY substitution model and YuleModel with 100 million iterations, every 1000 trees subsampling and discarding 25% of samples as burn-in. The log file from BEAST was analyzed using Tracer v1.7.2 [29] to verify all the parameter has effective sampling size (ESS) above 200 and tree was visualized/edited using Figtree v 1.4.4 [30]. Further, the mean genetic distance between our 13 unknown *Campylobacter spp*. samples and other *Campylobacter spp*. found in the neighboring clades from the phylogenetic analysis, were estimated through the Kimura 2-parameter (K2P) distance measure using MEGA 11 v11.0.10 [31].

## Results

Out of 67 fecal samples, 64 (95.5%) had detectable *Campylobacter*. Some soil (n=1) and water (n=1) samples were also contaminated with the bacteria (Table 1).

**Table 1:** *Campylobacter* in fecal samples of rhesus macaque, soil and water from two sites in Kathmandu.

|  | Total number of sample | Number (%) *Campylobacter* spp. present |
| --- | --- | --- |
| Rhesus Macaque Fecal | 67 | 64 (95.5%) |
| Water | 5 | 1 (20%) |
| Soil | 11 | 1 (0.9%) |

### Sequencing and Phylogenetic Analysis

Out of the 64 samples, only 13 provided acceptable quality of 16S DNA sequences, which was used to conduct phylogenetic analysis to resolve taxonomy of the *Campylobacter* isolates.

The topology of the phylogenetic tree constructed showed that different species of *Campylobacter spp*. clustering together in distinct clades. The isolates having same host species grouped together in a same clade with some exceptions. Our isolates clustered into distinct two clades identified-Clade-1 and Clade-2 (Figure 3). The clade-2 samples (PE005, PB002 and SA003) clustered together with *C. upsaliensis, C. vulpis, C. helveticus, C. troglodytes*. Whereas clade-1 (SA002, SA004 – SA009, SA011, SB008 & PD003) samples clustered together in a monophyletic clade comprising of a recently discovered species-*Candidatus Campylobacter infans*.

**Figure 1:**
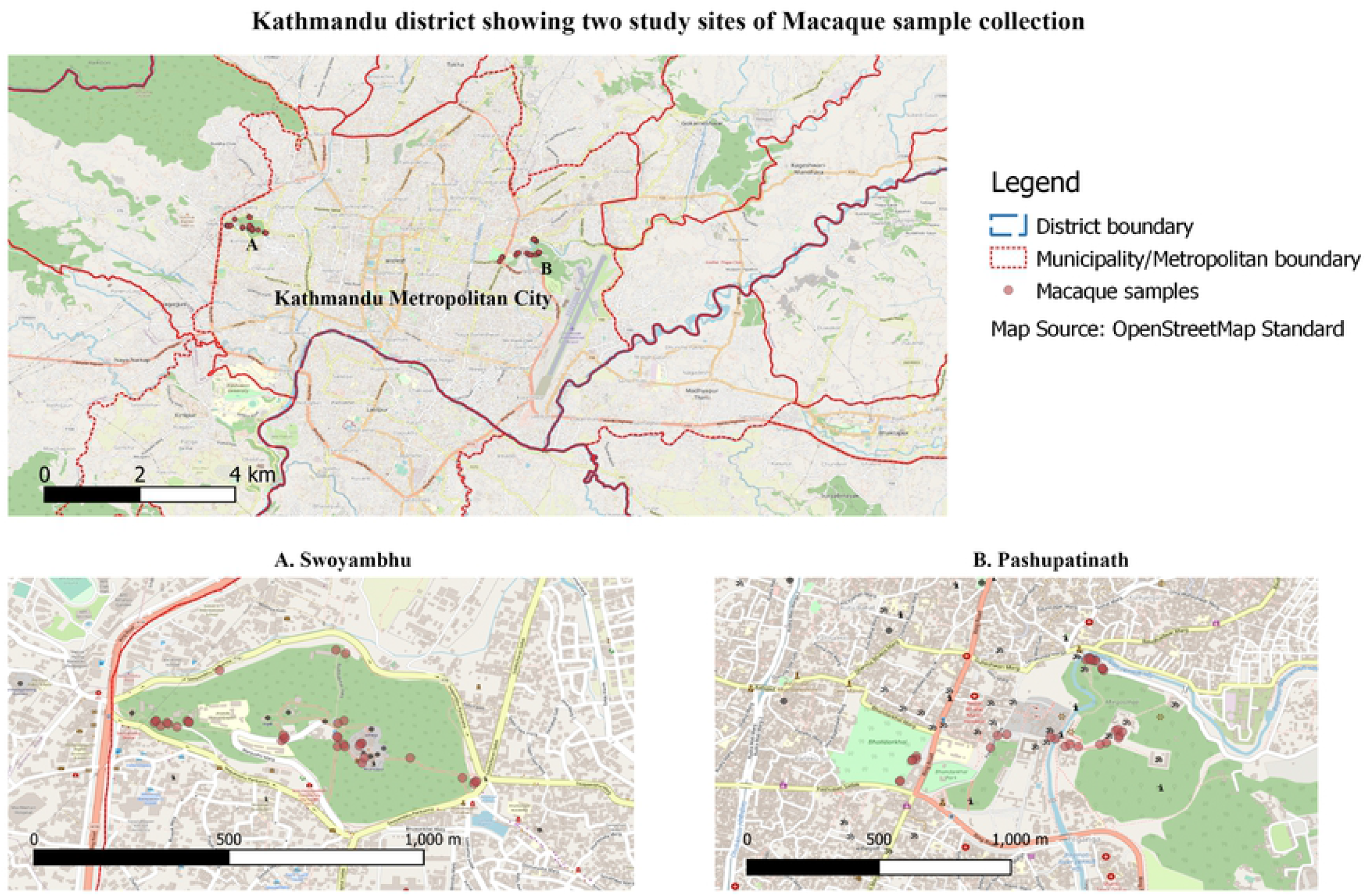
Rhesus macaque diarrheal outbreak sites (A. Swoyambhu B. Pashupatinath) located within Kathmandu Metropolitan city, Nepal (Image was created using QGIS).

**Figure 2:**
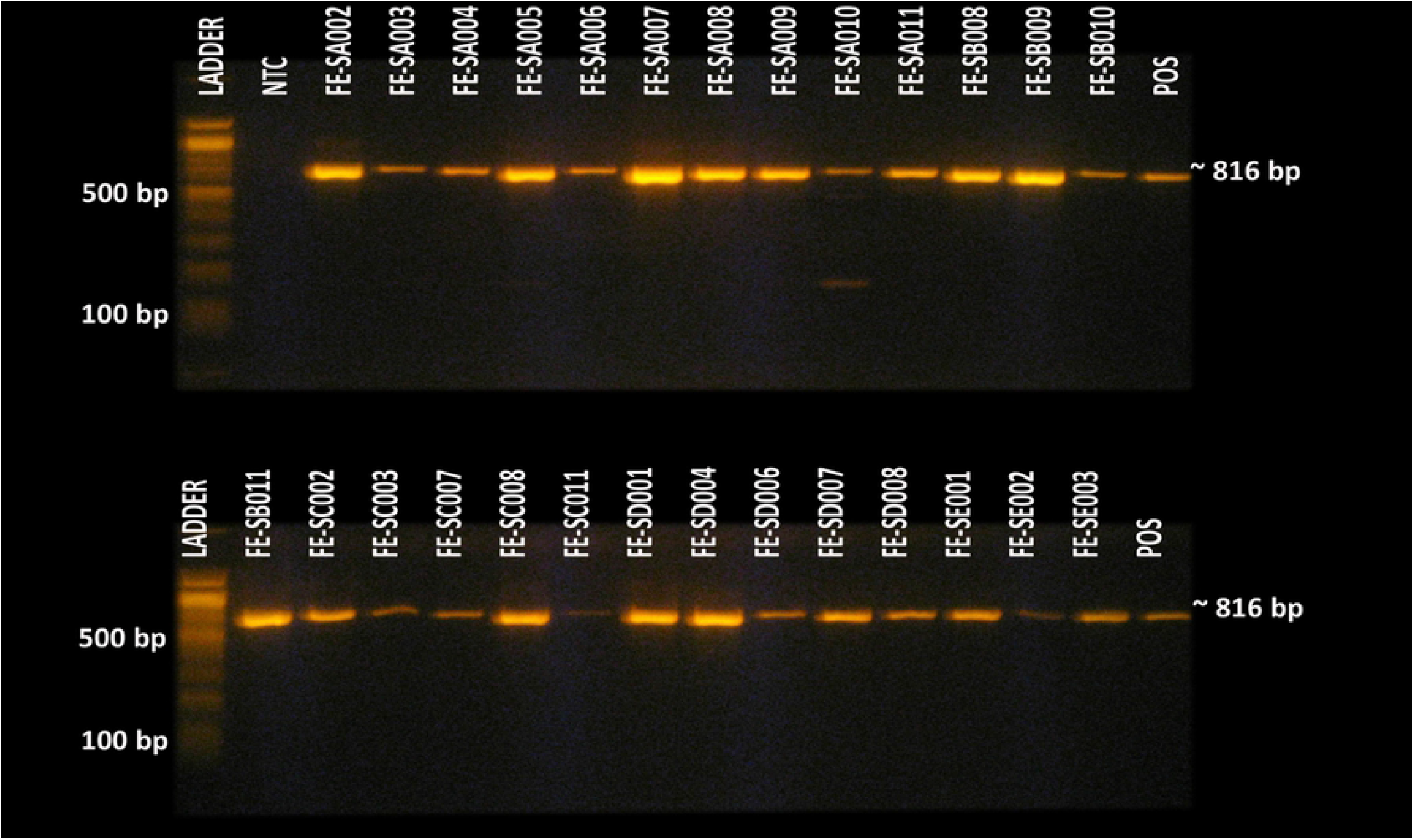
Detection of Campylobacter sp. in rhesus macaque fecal samples using PCR.

**Figure 3:**
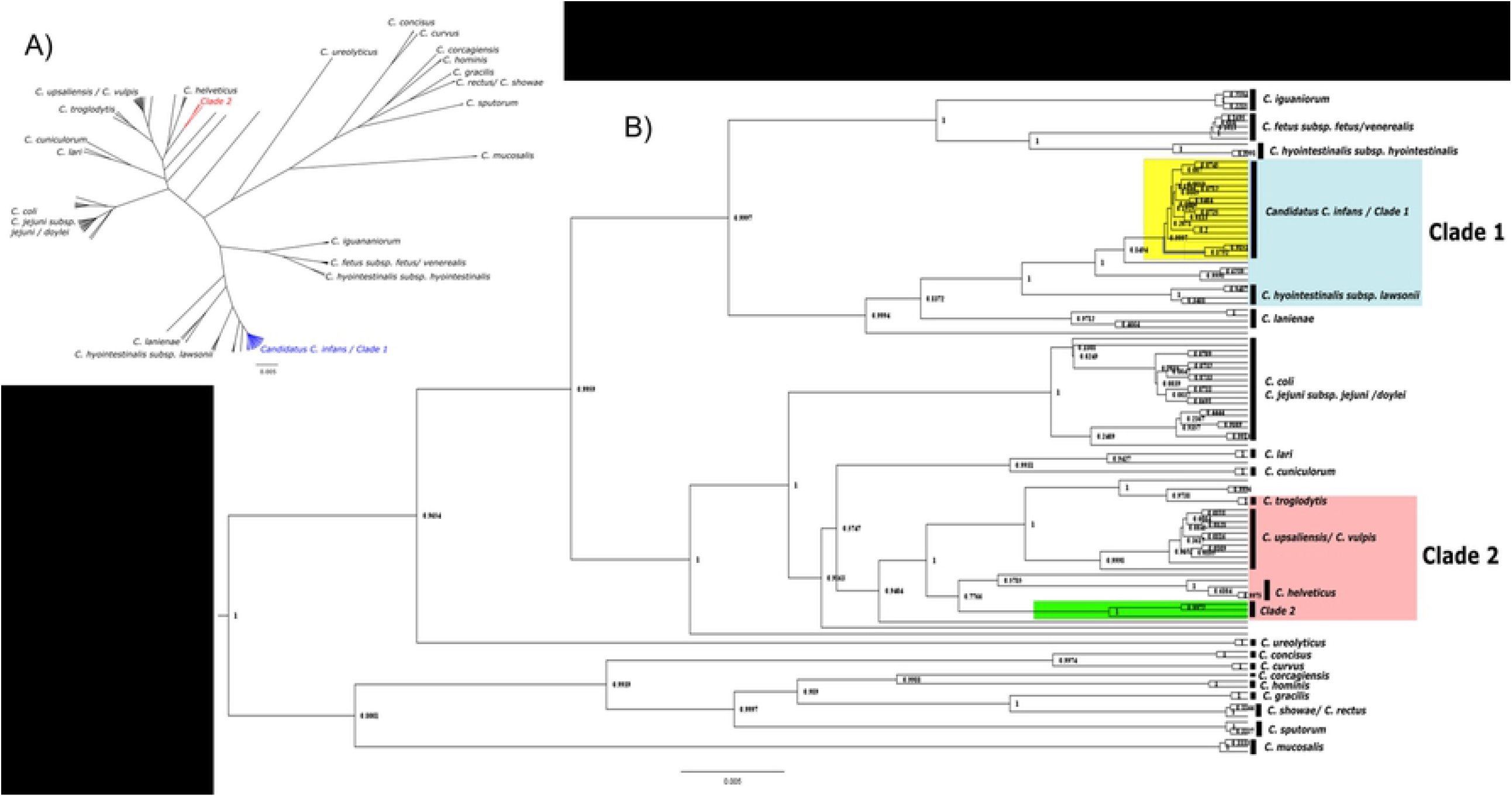
Phylogenetic tree of *Campylobacter spp*. showing all the detected and reference sequences [constructed using Bayesian inference (BEAST v 2.6.4) and visualized/edited using Figtree v 1.4.4]. A) Cladogram of *Campylobacter spp*. B) Phylogenetic tree constructed using detected and reference *Campylobacter spp*.

The Kimura 2-paramater (K2P) pairwise distance between clade-1 isolates and the recently discovered species of *Candidatus Campylobacter infans* (Genbank acc: CP049075) was 0.002287 after averaging the pairwise distances (supplementary figure 2). Similarly, the average K2P pairwise distance between clade-1 isolates with an isolate of *C. hyointestinalis subsp. lawsonii* (Genbank acc: HQ628645) found in the neighboring clade to the isolates, was calculated to be 0.0206. The average K2P pairwise distance of the clade-2 isolates was compared with a representative isolate of monophyletic clades of *C. upsaliensis* (Genbank acc: AF550642), *C. troglodytes* (Genbank acc: EU559331) and *C. helveticus* (Genbank acc: DQ174161) and was calculated to be 0.033, 0.0285, and 0.0267, respectively (supplementary figure 3).

## Discussion

Zoonotic pathogens are one of the most common sources of emerging diseases [32]. Campylobacter is considered to be one of the most prevalent zoonotic pathogens [33] that might be contributing to a broader antimicrobial resistance, especially in low and middle-income countries [34,35]. Nepal has very high burden of *Campylobacter* infections [5,36,37] and it is implicated for causing diarrhea in some of the international travelers; it is also known to cause acute gastroenteritis in children in Nepal [38–40]. Rhesus macaques (monkeys) have close-interactions with humans and other domestic animals, including dogs, in some areas of densely populated Kathmandu. The risk of zoonosis and reverse zoonosis of disease between monkeys and humans are high [33].

The detection and diagnosis of *Campylobacter* infection has been challenging due to inefficiency in widely used culture-based detection[41]. However, availability of molecular based diagnostic techniques have proved effective in detecting bacterial species that are normally difficult to culture [42,43]. This might be the first study in Nepal where *Campylobacter* is directly detected in macaque feces. The C*ampylobacter* was detected in water and soil samples as well, which could increase pathogen spillover possibility amongst various species intermingling at the sites [44].

The phylogenetic analysis of 16S rRNA gene sequences of *Campylobacter* showed presence of two clades of the bacteria (Fig. 3). The phylogenetic tree in our study also revealed more or less identical (Supplementary Fig. 4) structure of bacteria clades as found in the study published by Wilkinson D. A., et.al. (2018) [45]. That study used more elaborate whole genome sequence data, and the fact that our study also produced similar *Campylobacter* clade structure using only 16S rRNA fragment sequence data, validates the utility and accuracy of our technique.

The clade-1 isolates clustered together with the newly reported species of *Campylobacter* (*Candidatus C. infans*), though neighboring clades consisted of other species as well (Fig. 3). However, these isolates of other species in clade-1 could have been misidentified by the authors, as the K2P distances of isolates found in clade-1 have close distances to *Candidatus C. infans*, (Supplementary Figure 2) inferring that all isolates in clade-1 could possibly be identified as this newly discovered species. Previous cases of *Candidatus C. infans* have been isolated from infants from sub-Saharan Africa and South Asia, and from a male (human) in the Netherlands [46]. The *Campylobacter spp*. isolated in this study all originated from monkey fecal samples, which raises a serious concern over zoonotic capability of these particular isolates [47,48]. Furthermore, an isolate from India found in clade-1 was isolated from sheep alluding to strong evidence to support zoonotic plasticity.

Clade-2 contained the three remaining sequences from this study. The clustering of C. *helveticus, C. upsaliensis* and *C. troglodytes* in a nearby clades could also be observed as sister clades to clade-2 isolates. Our three isolates formed separate monophyletic clades (Fig. 3) within clade-2 indicating the findings of a probable new species or sub-species of the *Campylobacter*, which was further supported by the K2P pairwise distance with the closest distance of 0.0267 being with *C. helveticus*, which is similar to the values of clade-1 isolates against *C. hyointestinalis subsp. lawsonii* (HQ628645). However, further investigation with a longer 16S rRNA gene fragment or whole genome sequences may be required to properly verify the inferred result.

Samples were collected from two different sites separated by dense human settlements-almost making them wooded islands within the urban landscape. The *Campylobacter* isolates from both the sites clustered together in both clade-1 and clade-2 (Fig. 3). This result suggests there might be some complex interactions taking place between these two animal populations. Since monkeys from these two sites rarely intermingle, the disease spread might be limited as well. However, humans might help spread *Campylobacter* between these two populations of monkeys.

*C. hyointestinalis* is a commensal organism of pigs whereas *C. helveticus* is commensal in dogs and cats [45]. *Campylobacter* can potentially infect monkeys from these animals, and/or through humans as intermediate hosts. Considering other species such as birds, cats and dogs are also interacting closely with the monkeys at these two sites, the *Campylobacter* reservoir, spillover and transmission are truly playing out in One Health dynamics. Hence, this study highlights the importance of One Health approach to understand and prevent emerging, re-emerging and prevalent infectious diseases.

Diarrheal diseases are one of the most devastating public health concerns in Nepal, especially in a densely populated metropolitan cities like Kathmandu. *Campylobacter* might be one of the important contributing pathogens in diarrheal outbreaks-both in humans and animals (monkeys). We hope that our study will draw attention to this problem and help public health experts in formulating a plan to cure and prevent this kind of outbreak in macaques, thereby, preventing spillover to humans.

## Data Availability

DNA sequences are available in the NCBI Genbank with accession number MZ06810 to MZ068112. All the data are included in manuscript including supplementary information.

## Acknowledgement

We would like to thank the Pashupati Area Development Trust and the Federation of Swoyambhu Management and Conservation for providing the permit and facilitating this research. We would also like to thank PREDICT Consortium for their assistance during various phases of the project. We would also like to show our gratitude to the Metropolitan City of Kathmandu for helping us with the study. Finally, we thank all the staffs of Intrepid Nepal including Biswo Parakram Shrestha, Samita Raut, Dhiraj Puri for assisting in field sampling and lab experiments during the course of this study.

## Supplementary Information

**Supplementary Figure 1:** 16S rRNA based PCR detection of *Campylobacter sp*. from 27 Rhesus macaques fecal samples

**Supplementary Table 1:** Location of collected diarhhea fecal samples of Rhesus macaque

**Supplementary Table 2:** Reference sequences 16S rRNA used in this study

**Supplementary Figure 2:** Kimura-2 parameter pairwise distance calculation of clade-1 isolates compared to isolates of *C. hyointestinalis subsp. lawsonii* and *Candidatus C. infans*

**Supplementary Figure 3:** Kimura-2 parameter pairwise distance calculation of clade-2 with isolates of *C. vulpis, C. upsaliensis, C. troglodytes & C. helveticus*

**Supplementary Figure 4:** Comparison of core genome alignment network for non-thermophilic *Campylobacter* species (right) from D. A. Wilkinson, et.al., (2018) and partial 16srRNA gene sequences obtained in this study. Phylogenetics prepared by using BEAST v 2.6.4 (left). Arrow indicates clade from same species (for ease of comparison).

## Reference

1. Andrade MCR, Gabeira SC de O, Abreu-Lopes D, Esteves WTC, Vilardo M de CB, Thomé J, et al. Circulation of Campylobacter spp. in rhesus monkeys (Macaca mulatta) held in captivity: a longitudinal study. Mem Inst Oswaldo Cruz. 2007;102: 53–57.

2. Komba EV. Human and Animal Thermophilic Campylobacter infections in East African countries: Epidemiology and Antibiogram. Biomed J Sci Tech Res. 2017;1. doi:10.26717/BJSTR.2017.01.000411

3. Weis AM, Storey DB, Taff CC, Townsend AK, Huang BC, Kong NT, et al. Genomic comparison of Campylobacter spp. and their potential for zoonotic transmission between birds, primates, and livestock. Appl Environ Microbiol. 2016;82: 7165–7175.

4. Devaux CA, Mediannikov O, Medkour H, Raoult D. Infectious disease risk across the growing human-non human primate interface: A review of the evidence. Front Public Heal. 2019; 305.

5. Acharya KP, Wilson RT. Antimicrobial resistance in Nepal. Front Med. 2019;6: 105.

6. Bhattarai D, Bhattarai N, Osti R. Prevalence of Thermophilic Campylobacter Isolated from Water Used in Slaughter House of Kathmandu and Ruphendehi District, Nepal. Int J Appl Sci Biotechnol. 2019;7: 75–80.

7. Gautam A, Neupane S, Kaphle K. Campylobacter Infection and its Sensitivity in Retail Pork. Int J Appl Sci Biotechnol. 2020;8: 132–139.

8. Neupane R, Kaphle K. Bacteriological quality of poultry meat in Nepal. Int J Vet Sci. 2019.

9. Abebe E, Gugsa G, Ahmed M. Review on major food-borne zoonotic bacterial pathogens. J Trop Med. 2020;2020.

10. Pathak A, Kaphle K. Dog: A Friendly Pathway to Zoonoses. Nepal Vet J. 2019;36: 170–177.

11. Adhikari R, Bagale KB. Risk of Zoonoses among livestock farmers in Nepal. J Heal Promot. 2019;7: 99–110.

12. Misawa N, Shinohara S, Satoh H, Itoh H, Shinohara K, Shimomura K, et al. Isolation of Campylobacter species from zoo animals and polymerase chain reaction-based random amplified polymorphism DNA analysis. Vet Microbiol. 2000;71: 59–68. doi:10.1016/S0378-1135(99)00156-X

13. Ngotho M, Ngure RM, Kamau DM, Kagira JM, Gichuki C, Farah IO, et al. A fatal outbreak of Campylobacter jejuni enteritis in a colony of vervet monkeys in Kenya. 2017.

14. Keita MB, Hamad I, Bittar F. Looking in apes as a source of human pathogens. Microb Pathog. 2014;77: 149–154. doi:10.1016/J.MICPATH.2014.09.003

15. Rhoades N, Barr T, Hendrickson S, Prongay K, Haertel A, Gill L, et al. Maturation of the infant rhesus macaque gut microbiome and its role in the development of diarrheal disease. Genome Biol. 2019;20: 1–16.

16. Clayton JB, Danzeisen JL, Johnson TJ, Trent AM, Hayer SS, Murphy T, et al. Characterization of Campylobacter jejuni, Campylobacter upsaliensis, and a novel Campylobacter sp. in a captive non-human primate zoological collection. J Med Primatol. 2018/12/12. 2019;48: 114–122. doi:10.1111/jmp.12393

17. Sowerby N. Identification and genotyping of Campylobacter spp. strains isolated from a captive wildlife population in New Zealand. Auckland University of Technology. 2015.

18. Tegner C, Sunil-Chandra NP, Wijesooriya W, Perera BV, Hansson I, Fahlman Å. Detection, Identification, and Antimicrobial Susceptibility of Campylobacter spp. and Salmonella spp. from Free-ranging Nonhuman Primates in Sri Lanka. J Wildl Dis. 2019;55: 879–884.

19. Adams N, Dhimal M, Mathews S, Iyer V, Murtugudde R, Liang X-Z, et al. El Niño Southern Oscillation, monsoon anomaly, and childhood diarrheal disease morbidity in Nepal. PNAS Nexus. 2022;1: pgac032.

20. Chalise MK. Primate Census in Kathmandu and West Parts of Nepal. J Nat Hist Mus. 2008;23: 60–64. doi:10.3126/JNHM.V23I0.1840

21. Chalise MK. Fragmented Primate Population of Nepal. Primates Fragm Complex Resil. 2013; 329–356. doi:10.1007/978-1-4614-8839-2_22

22. Baral K. Conservation and threat to selected monkey species in Nepal compared to selected species in Tanzania. Norwegian University of Life Sciences. 2014.

23. Linton D, Owen RJ, Stanley J. Rapid identification by PCR of the genus Campylobacter and of five Campylobacter species enteropathogenic for man and animals. Res Microbiol. 1996;147: 707–718. doi:10.1016/S0923-2508(97)85118-2

24. Bouckaert R, Vaughan TG, Barido-Sottani J, Duchêne S, Fourment M, Gavryushkina A, et al. BEAST 2.5: An advanced software platform for Bayesian evolutionary analysis. PLoS Comput Biol. 2019;15: e1006650. doi:10.1371/journal.pcbi.1006650

25. Gorkiewicz G, Feierl G, Zechner R, Zechner EL. Transmission of Campylobacter hyointestinalis from a pig to a human. J Clin Microbiol. 2002;40: 2601–2605. doi:10.1128/JCM.40.7.2601-2605.2002/ASSET/A946BEDE-3BB4-4777-A449-8F5DCD18D429/ASSETS/GRAPHIC/JM0721442001.JPEG

26. Edgar RC. MUSCLE: multiple sequence alignment with high accuracy and high throughput. Nucleic Acids Res. 2004;32: 1792–1797. doi:10.1093/NAR/GKH340

27. Larsson A. AliView: a fast and lightweight alignment viewer and editor for large datasets. Bioinformatics. 2014;30: 3276–3278. doi:10.1093/bioinformatics/btu531

28. Bouckaert RR, Drummond AJ. bModelTest: Bayesian phylogenetic site model averaging and model comparison. BMC Evol Biol. 2017;17: 1–11. doi:10.1186/S12862-017-0890-6/FIGURES/6

29. Rambaut A, Drummond AJ, Xie D, Baele G, Suchard MA. Posterior Summarization in Bayesian Phylogenetics Using Tracer 1.7. Syst Biol. 2018;67: 901–904. doi:10.1093/SYSBIO/SYY032

30. Rambaut A. Figtree v1.4.4.

31. Tamura K, Stecher G, Kumar S. MEGA11: Molecular Evolutionary Genetics Analysis Version 11. Mol Biol Evol. 2021;38: 3022–3027. doi:10.1093/MOLBEV/MSAB120

32. Magouras I, Brookes VJ, Jori F, Martin A, Pfeiffer DU, Dürr S. Emerging zoonotic diseases: Should we rethink the animal–human interface? Front Vet Sci. 2020; 748.

33. Masila NM, Ross KE, Gardner MG, Whiley H. Zoonotic and Public Health Implications of Campylobacter Species and Squamates (Lizards, Snakes and Amphisbaenians). Pathogens. 2020;9: 799.

34. Hlashwayo DF, Sigaúque B, Bila CG. Epidemiology and antimicrobial resistance of Campylobacter spp. in animals in Sub-Saharan Africa: A systematic review. Heliyon. 2020;6: e03537.

35. Kaakoush NO, Castaño-Rodríguez N, Mitchell HM, Man SM. Global epidemiology of Campylobacter infection. Clin Microbiol Rev. 2015;28: 687–720.

36. Ghimire L, Singh DK, Basnet HB, Bhattarai RK, Dhakal S, Sharma B. Prevalence, antibiogram and risk factors of thermophilic campylobacter spp. in dressed porcine carcass of Chitwan, Nepal. BMC Microbiol. 2014;14: 85. doi:10.1186/1471-2180-14-85

37. Gautam A, Neupane S, Kaphle K. Campylobacter Infection and its Sensitivity in Retail Pork. Int J Appl Sci Biotechnol. 2020;8: 132–139. doi:10.3126/IJASBT.V8I2.29587

38. Bhattarai V, Sharma S, Rijal KR, Banjara MR. Co-infection with Campylobacter and rotavirus in less than 5 year old children with acute gastroenteritis in Nepal during 2017-2018. BMC Pediatr. 2020;20: 1–8. doi:10.1186/S12887-020-1966-9/FIGURES/2

39. Murphy H, Bodhidatta L, Sornsakrin S, Khadka B, Pokhrel A, Shakya S, et al. Traveler’s diarrhea in Nepal-changes in etiology and antimicrobial resistance. J Travel Med. 2019;26. doi:10.1093/JTM/TAZ054

40. Pandey P, Bodhidatta L, Lewis M, Murphy H, Shlim DR, Cave W, et al. Travelers’ diarrhea in Nepal: an update on the pathogens and antibiotic resistance. J Travel Med. 2011;18: 102–108. doi:10.1111/J.1708-8305.2010.00475.X

41. CDC. Centers for disease control and prevention, national center for emerging and zoonotic infectious diseases (NCEZID), Division of Foodborne, Waterborne, and Environmental Diseases (DFWED). 2012.

42. do Nascimento Veras H, da Silva Quetz J, Lima IFN, Rodrigues TS, Havt A, Rey LC, et al. Combination of different methods for detection of Campylobacter spp. in young children with moderate to severe diarrhea. J Microbiol Methods. 2016;128: 7–9.

43. Ricke SC, Feye KM, Chaney WE, Shi Z, Pavlidis H, Yang Y. Developments in rapid detection methods for the detection of foodborne campylobacter in the United States. Front Microbiol. 2019; 3280.

44. Facciolà A, Riso R, Avventuroso E, Visalli G, Delia SA, Laganà P. Campylobacter: from microbiology to prevention. J Prev Med Hyg. 2017;58: E79. Available: /pmc/articles/PMC5584092/

45. Wilkinson DA, O’Donnell AJ, Akhter RN, Fayaz A, MacK HJ, Rogers LE, et al. Updating the genomic taxonomy and epidemiology of Campylobacter hyointestinalis. Sci Reports 2018 81. 2018;8: 1–12. doi:10.1038/s41598-018-20889-x

46. Flipse J, Duim B, Wallinga JA, de Wijkerslooth LRH, Graaf-van Bloois L van der, Timmerman AJ, et al. A Case of Persistent Diarrhea in a Man with the Molecular Detection of Various Campylobacter species and the First Isolation of candidatus Campylobacter infans. Pathogens. 2020;9: 1003.

47. Delahoy MJ, Wodnik B, McAliley L, Penakalapati G, Swarthout J, Freeman MC, et al. Pathogens transmitted in animal feces in low-and middle-income countries. Int J Hyg Environ Health. 2018;221: 661–676.

48. Himsworth CG, Skinner S, Chaban B, Jenkins E, Wagner BA, Harms NJ, et al. Multiple zoonotic pathogens identified in canine feces collected from a remote Canadian indigenous community. Am J Trop Med Hyg. 2010;83: 338.

49. Ahmed W, Hodgers L, Sidhu JPS, Toze S. Fecal indicators and zoonotic pathogens in household drinking water taps fed from rainwater tanks in Southeast Queensland, Australia. Appl Environ Microbiol. 2012;78: 219–226.

50. Ferreira V, Magalhães R, Teixeira P, Castro PML, Calheiros CSC. Occurrence of Fecal Bacteria and Zoonotic Pathogens in Different Water Bodies: Supporting Water Quality Management. Water. 2022;14: 780.

